# *unc-37/*Groucho and *lsy-22/*AES repress Wnt target genes in *C. elegans* asymmetric cell divisions

**DOI:** 10.1101/2022.01.10.475695

**Authors:** Kimberly N. Bekas, Bryan T. Phillips

## Abstract

Asymmetric cell division (ACD) is a fundamental mechanism of cell fate specification and adult tissue homeostasis. In *C. elegans*, the Wnt/β-catenin asymmetry (WβA) pathway regulates ACDs throughout embryonic and larval development. Under control of Wnt ligand-induced polarity, the transcription factor POP-1/TCF functions with the coactivator SYS-1/β-catenin to activate gene expression in the signaled cell or, in absence of the coactivator, to repress Wnt target genes in the unsignaled daughter cell. To date, investigation of Groucho function in WβA is lacking, and the function of LSY-22/AES has only been evaluated in *C. elegans* neurons. Further, conflicting evidence shows TCF utilizing Groucho-mediated repression may be either aided or repressed by AES addition. Here we demonstrate a genetic interaction between Groucho corepressors and POP-1/TCF in the distal tip cells (DTCs), seam cells (SCs) and embryonic endoderm development. In the DTCs, signaled cell fate increases after individual and double Groucho loss of function, representing the first demonstration of Groucho function in wildtype WβA ACDs. Further, WβA target gene misexpression occurs more frequently than DTC fate changes, suggesting derepression generates an intermediate cell fate. In the SCs, loss of UNC-37/Groucho or LSY-22/AES in a POP-1/TCF hypomorphic background enhances SC expansion and target gene misregulation. Moreover, while POP-1/TCF depletion in *lsy-22/AES* nulls yielded an expected increase in SCs we observed a surprising SC decrease in *unc-37/Groucho* nulls subjected to POP-1/TCF depletion. This phenotype correlates with UNC-37/Groucho regulation of *pop-1/tcf* expression since POP-1/TCF levels are increased in *unc-37/Groucho* null SCs. Lastly, Groucho functions in embryonic endoderm development since we observe ectopic endoderm transgene expression in *unc-37/Groucho* and *lsy-22/AES* knockdown in a HDA-1 background. Together, these data indicate Groucho-mediated modulation of cell fate via regulation of POP-1/TCF repression is widespread in WβA ACDs and suggests a novel role of LSY-22/AES as a *bona fide* TCF repressor.

## Introduction

The development of multicellular organisms necessitates generating a variety of specialized cells from a single fertilized egg (Conklin, 1905). One way this is accomplished is via asymmetric cell divisions (ACDs), where a dividing mother cell asymmetrically localizes cytoplasmic determinants to opposing ends of a cell (Conklin, 1905; Rhyu, Jan, & Jan, 1994). This process gives rise to two daughter cells that have distinct cell fates at birth, contributing to the development of distinct tissues in subsequent lineages. ACD is an essential process for generating cell diversity during development and maintaining adult tissue homeostasis via stem cell specification (Horvitz & Herskowitz, 1992; Neumuller & Knoblich, 2009). Importantly, when the pathways regulating ACD go awry, tumorigenesis and forms of cancer result (Blaydon et al., 2006; Parma et al., 2006).

The Wnt/β-catenin asymmetry pathway (WβA) is a well-established critical regulator of ACDs (Schneider & Bowerman, 2007). In the signaled daughter cell resulting from a Wnt-signaled ACD, the Wnt ligand binds to the Wnt receptor, composed of Frizzled (Fz) and LRP6, resulting in destabilization of the destruction complex (He, Semenov, Tamai, & Zeng, 2004; Logan & Nusse, 2004). With an ineffective destruction complex, the coactivator, β-catenin, accumulates in the cytoplasm and eventually translocates into the nucleus. Once in the nucleus, the β-catenin coactivator interacts with the transcription factor TCF to activate Wnt target gene expression. The WβA pathway uses an external Wnt gradient to asymmetrically regulate the Wnt signaling status between the two daughter cells. In the unsignaled daughter cell resulting from a Wnt-polarized ACD, lack of Wnt ligand allows for the stabilized cytoplasmic destruction complex to phosphorylate β-catenin, ultimately leading to its proteasomal degradation (He et al., 2004; Kimelman & Xu, 2006; Logan & Nusse, 2004). Without a nuclear coactivator, TCF remains bound to the DNA where it is thought to function to turn off Wnt target genes. While typically Wnt target gene repression in the unsignaled daughter is known to result from TCF-mediated repression, the repressive mechanism is not well characterized.

In canonical Wnt signaling, TCF is known to interact with the Groucho/Transducin-like Enhancer of Split family of corepressors (Daniels & Weis, 2005). Therefore, one possible mechanism for TCF target gene repression in the WβA pathway is interaction with Groucho corepressors. Thus, investigation to determine if Groucho-mediated functions in in the WβA pathway is warranted.

The Groucho protein family are corepressors, meaning they cannot directly bind DNA but instead bind to transcription factors to confer repression. Transcription factor binding occurs via one of two highly conserved domains: the N-terminal, glutamine-rich Q domain or the C-terminal WD-repeat domain (Chen, Nguyen, & Courey, 1998; Miyasaka, Choudhury, Hou, & Li, 1993; Pinto & Lobe, 1996; Song, Hasson, Paroush, & Courey, 2004). In fact, the Groucho family is broken down in two subfamilies based on the presence of one or both highly conserved domains. Family members containing both the Q domain and the WD-repeat domain are full-length Groucho proteins belonging to the Groucho subfamily. Truncated family members containing the Q domain but lacking the WD-repeat domain belong to the amino-terminal enhancer of split (AES) subfamily, sometimes referred to as short Grouchos. Notably, Groucho and AES binding to TCF occurs via the Q domain (Chen et al., 1998; Miyasaka et al., 1993; Pinto & Lobe, 1996; Song et al., 2004). Since both Groucho and AES family members contain the Q domain, it is possible for both Groucho and AES proteins to interact with TCF. Despite this potential for interaction, the role of Groucho and AES function in WβA-signaled ACDs has yet to be fully interrogated.

After transcription factor binding, the mechanism for Groucho-mediated repression is generally broken into two categories. First, yeast genetic and biochemical studies demonstrated a Groucho-related corepressor, Tup1, interacts with the RNA Pol II subunits and the Mediator complex (Berk, 1999; Kornberg, 2005; Malik & Roeder, 2010). Further studies of the *C. elegans* Groucho homolog *unc-37/Groucho* indicate that it may interact with Mediator in the adult male seam cells that give rise to the mail tail rays (Zhang & Emmons, 2002). Taken together, these studies exemplify that Groucho proteins may confer repression via blockage of transcriptional machinery. Secondly, Groucho-mediated repression is known to change chromosomal structure; Groucho proteins in yeast, flies, and mammals are capable of binding to all four core histones via their N-terminal tails (Edmondson and Roth 1998, Edmonson et al 1996, Flores-saaib and Courey 2000; Palparti et al 1997). Given these two options, it is possible that Groucho-mediated repression may occur in a variety of potentially context dependent manners.

*Caenorhabditis elegans* is well suited for investigating the role of Groucho proteins in WβA regulated ACDs because there is only one Groucho protein, known as *unc-37/Groucho*, and one AES protein, known as *lsy-22/AES*. This allows the study of the roles of each subfamily member without redundancy precluding analysis. Previous work on *unc-37/Groucho* and *lsy-22/AES* shows they are widely, if not constitutively, expressed from the 4-cell embryonic stage through larval stage 4 (Flowers et al., 2010; Pflugrad, Meir, Barnes, & Miller, 1997). *C. elegans* Groucho and AES homologs also regulate proper cell fate decisions. In neuronal development, UNC-37/Groucho and LSY-22/AES work together with the transcription factor COG-1 for proper ASER cell fate and with UNC-4 for proper VA/VB cell fate (Flowers et al., 2010; Winnier et al., 1999). Consistent with conserved family member function, UNC-37/Groucho is known to physically interact with POP-1, the *C. elegans* TCF homolog (Calvo et al., 2001). Given UNC-37/Groucho and LSY-22/AES expression, function in cell fate determination, and POP-1/TCF interaction, it is possible these general corepressors play a role in WβA cell fate decisions.

Further, truncated AES Groucho family members classically function to sequester, and thereby negatively regulate, full-length Groucho protein (Bajoghli, Aghaallaei, Soroldoni, & Czerny, 2007; Beagle & Johnson, 2010; Li, 2000) This model is challenged by findings in *C. elegans* neuronal development. Here, loss of either *unc-37/Groucho* or *lsy-22/AES* resulted in the same phenotypic changes in neuronal cell fate via interaction with transcription factors (Flowers et al., 2010). Thus, evidence for AES function as both a negative regulator and cooperative partner of Groucho exists. Further, both *unc-37/Groucho* and *lsy-22/AES* contain the required domain for TCF binding. Therefore, we investigated both *C. elegans* Grouchos in different developmental contexts to determine the composition and function of the corepressor complex.

The *C. elegans* model organism is well suited to elucidate the corepressor function in different contexts as it contains three well characterized WβA-signaled ACDs: the distal tip cells (DTCs), the seam stem cells (SCs) and embryonic endoderm specification. Of these three, the DTCs are thought to be an activation dominant tissue, since loss of the TCF coactivator SYS-1/β-catenin is results in a loss of DTC fate (Siegfried, Kidd, Chesney, & Kimble, 2004). Conversely, the SCs and endoderm are thought to be a repression dominant tissue since loss of SYS-1/β-catenin does not change SC or endoderm cell fate (Phillips, Kidd, King, Hardin, & Kimble, 2007). To date there has been only one study of *lsy-22/AES*, which did not investigate its role in WβA-signaled ACDs (Flowers et al., 2010). Therefore, a thorough investigation of the role of Groucho-mediated repression in Wnt-signaled ACDs is needed to better understand the mechanism of proper cell fate specification.

Here, we show that Groucho proteins promote repression of the DTC fate after ACD as *unc-37/Groucho* and *lsy-22/AES* mutants show DTC expansion, a phenotype which is enhanced by knocking down *lsy-22/AES* or *unc-37/Groucho*, respectively. In the SC ACD, we found Groucho loss can enhance an intermediate *pop-1(RNAi)* phenotype. Further we showed in the SCs, *unc-37/Groucho* knockdown increases POP-1/TCF levels, indicating UNC-37/Groucho regulates *pop-1/tcf* in this tissue. We also showed in the activation dominant DTCs, Wnt target gene derepression occurred more frequently than cell fate changes occurred. Conversely, in the repression dominant SCs, target gene misexpression mirrored cell fate changes resulting from Groucho loss. During embryonic endoderm development, we show that *lsy-22/AES* functions similarly to previously investigated *unc-37/Groucho* function. Specifically, knockdown of either *unc-37/Groucho* or *lsy-22/AES* in a HDA-1 compromised background results in ectopic endoderm transgene expression. Altogether, we determined that both *unc-37/Groucho*, and the understudied *lsy-22/AES* corepressor, function in both activation dominant and repression dominant of WβA-signaled ACDs.

## Materials and Methods

### Strain list

Strains were maintained using standard *C. elegans* methods (Brenner, 1974). The strains used are as follows:

1. **JK2868** [*qIs56*[Plag-2::GFP + unc-119(+)]]
2. **JK3001** [*pop-1(q645*) (III); qIs56(V)]
3. **BTP226** [*unc-37(tm4649)*(I)/ hT2[bli-4(e937) let-?(q782) *qIs48*](I;III); *qIs56*(V)]
4. **BTP230** [*lsy-22(ot244)*(I)/ hT2[bli-4(e937) let-?(q782) *qIs48*](I;III); *qIs56*(V)]
5. **JR667** [*unc-119*(e2498::Tc1) (III); *wIs51*[P_scm_::GFP + unc-119(+)] (V)]
6. **BTP227** [*rde-1(ne219)*(V); *uIwEx33*[pAW559(P_scm_::RDE-1), Pttx-3::Dsred]; *wIs51*(V)]
7. **BTP224** [*unc-37(tm4649)*(I)/ hT2[bli-4(e937) let-?(q782) *qIs48*](I;III); *wIs51*(V)]
8. **BTP223** [*lsy-22(ot244)*(I)/ hT2[bli-4(e937) let-?(q782) *qIs48*](I;III); *wIs51*(V)]
9. **BTP241** [*pop-1*(he335[eGFP::loxP::pop-1])(I); *ouIs21*[ajm-1p::mCherry]
10. **BTP232** [*unc-37(tm4649)*/hT2[bli-4(e937) let-?(q782) *qIs48*]; *qIs90*]
11. **BTP231** [*lsy-22(ot244)*/hT2 [bli-4(e937) let-?(q782) *qIs48*]; *qIs90*]
12. **RW11606** [unc-119(tm4063) (III); *stIs11606* [egl-18a::H1-wCherry + unc-119(+)]]
13. **BTP243** [*unc-37(tm4649)* (I); *pop-1(he335[eGFP::loxP::pop-1])*(I); *ouIs21*[ajm-1p::mCherry]
14. **TX691** [*unc-119(ed3)* (III); *teIs46*]
15. **BTP245** [*unc-37(tm4649);lsy-22(uIw1*[Q33STOP; C97T, G99A](I)/tmC20[unc-14(tmIs1219) dpy-5(tm9715)]; *teIs46*]
16. **BTP239** [*unc-37(tm4648)*(I)/hT2[bli-4(e937) let-?(q782) *qIs48*]; *teIs46*]
17. **BTP26** [*lsy-22(ot244)*(I)/tmC20[unc-14(tmIs1219) dpy-5(tm9715)]; *teIs46*]

#### RNAi

RNAi constructs were expressed form *E. coli* HT115(DE3) bacteria grown on plates containing 1mM IPTG to induce T7 convergent promotors (Timmons & Fire, 1998). All target genes were obtained from the Ahringer library except *unc-37* and *lacZ* which were cloned using target gene amplification of *C. elegans* cDNA and *E. coli* DNA, respectively. Simultaneous RNAi by bacterial mixing was carried out by growing two different RNAi bacterial strains to the same optical density and mixing volumes in the ratios indicated before plating. RNAi was induced by L1 synchronized feeding until imaging the appropriate developmental stage.

#### CRISPR/Cas9

CRISPR/Cas9 was used to create the double Groucho null *unc-37(tm4649);lsy-22*(*uIw1*[Q33STOP; C97T, G99A]). Heterozygous *unc-37(tm4649)* hermaphrodites were injected with a mix containing 17.6 μM TracRNA (IDT), 16.2 μM *lsy-22* cRNA (IDT), 6 μM *lsy-22* ssODN (IDT), 1.5 μM *dpy-10* cRNA (IDT), 0.5 μM *dpy-10* ssODN (IDT), and 14.35 μM Cas-9-NLS (Berkeley Macro lab) (Hefel & Smolikove, 2019). Mix was made by duplexing tracRNA, target cRNA and *dpy-10* cRNA at 95°C for 5 minutes and allowing to cool at room temperature for 5 minutes. After duplexing, Cas9 was added and allowed to complex at room temperature before adding *dpy-10* and target ssODN. Young adult worms were injected and isolated to individual plates the morning following injection. Plates were screened for rol or dpy phenotypes created by the *dpy-10* co-CRISPR marker adopted from (Paix, Schmidt, & Seydoux, 2016) and sequenced to identify strains with the proper target edit.

#### Image Processing and quantification

Images were obtained using Zeiss Axio Imager.D2 compound fluorescent scope and Zeiss Zen software. Raw GFP image data was exported to a JPEG format and analyzed using ImageJ software. Nuclei of seam cells were identified by DIC images and then the mean protein fluorescence was quantified using ImageJ and normalized. All statistical comparisons were performed using Graph Prism software. Image of extra DTCs was obtained using Leica CS SPE Confocal Microscope RGBV and represented as a full Z-projection using the LAS AF SPE Core Software.

## Results

### UNC-37/Groucho and LSY-22/AES are required for proper distal tip cell fate

To determine if Groucho proteins are required in WβA-signaled ACDs, we began to investigate the DTCs. The DTCs comprise the adult worm germline stem cell niche. They are derived from somatic gonadal precursors Z1 and Z4 that divide asymmetrically twice in the first larval stage (L1) to give rise to the two terminally differentiated DTCs ( Figure 1A). Studies of Wnt signaling in the DTCs demonstrate loss of SYS-1/β-catenin decreases DTC fate (Figure 1B) (Siegfried et al., 2004). Since the coactivator is required for proper cell fate, the DTCs are considered an activation dominant tissue. Despite this categorization, we wanted to investigate the potential contribution of Groucho corepressors to DTC determination

**Figure 1:**
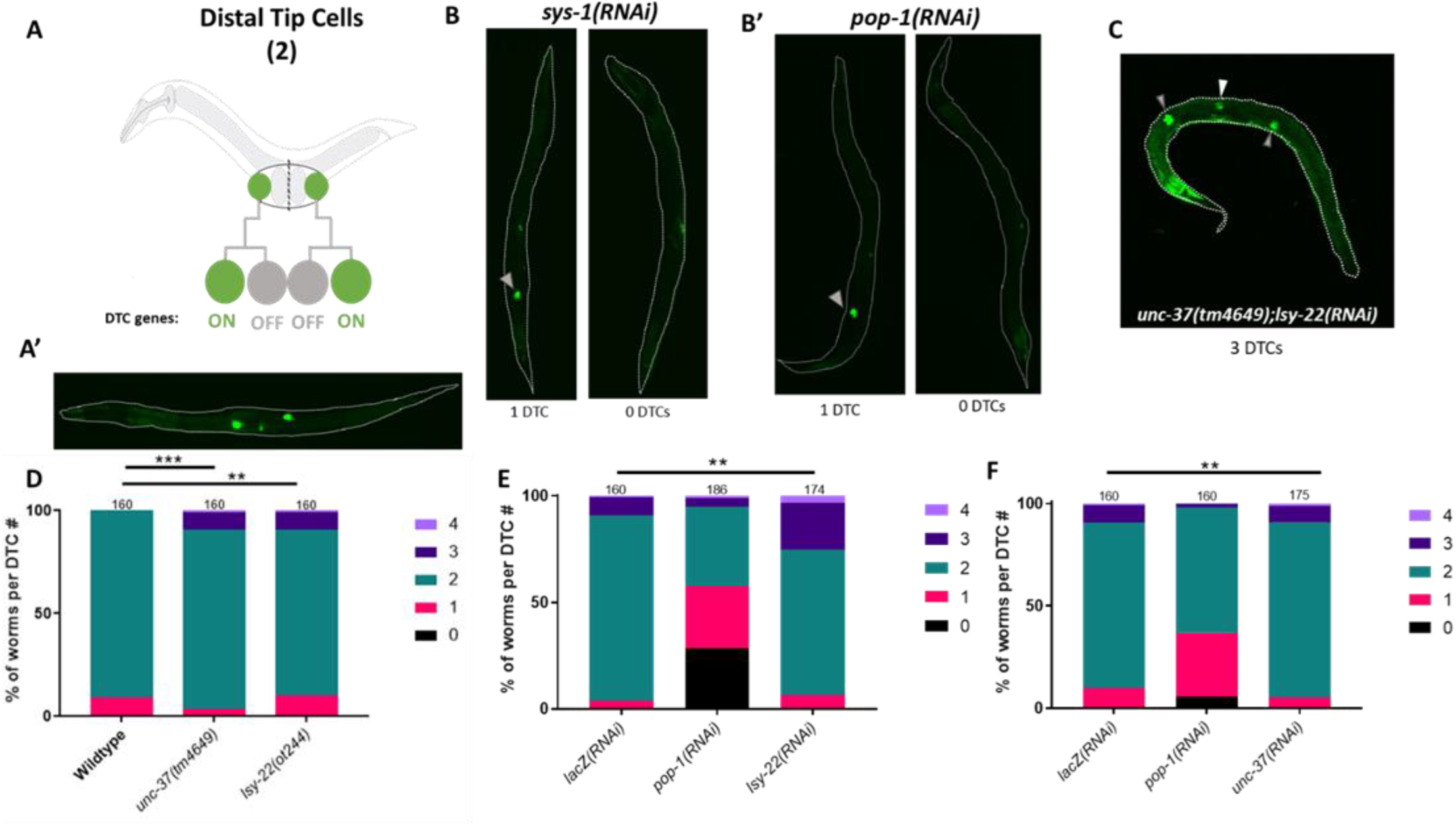
UNC-37/Groucho and LSY-22/AES affect distal tip cell fate. A) Graphic of the DTC lineage in a wildtype worm and A’) representative image of DTCs with *qIs56*[P_lag-2_::GFP]. B) Representative images of DTC loss resulting from *sys-1/β-catenin* or B’) *pop-1/tcf* knockdown. Endogenous DTCs are denoted by the gray arrowhead. C) Representative image of *unc-37/Groucho* null with one ectopic DTC denoted by the white arrowhead. Endogenous DTCs are denoted by the gray arrowheads. Image taken on Leica CS SPE Confocal Microscope RGBV shown as a full Z-projection using LAS AF SPE Core Software. D) Corepressor nulls increase DTC fate. E) *lsy-22/AES* knockdown enhances DTC fate gains in *unc-37/Groucho* null. F) *unc-37/Groucho* knockdown enhances DTC fate gains in *lsy-22/AES* null. N value listed above each bar. Colors correlate to the number of DTCs in an individual worm. *-p≤0.5, **-p<0.01, ***-p<0.001, ****-p≤ 0.0001 via unpaired t-Test.

First, we assessed the ability of *unc-37(RNAi)* or *lsy-22(RNAi)* individually, and together, to affect DTC fate in worms with GFP-marked DTCs (*qIs56*[P_*lag-2*_::GFP]). Neither individual nor double Groucho knockdown affected DTC fate in a wildtype background (Supplemental Figure 1A). However, in a *pop-1(q645)* mutant background, where POP-1/TCF binding to its coactivator SYS-1/β-catenin is impaired, we found individual Groucho knockdown mildly suppressed DTC loss (Phillips et al., 2007) (Supplemental Figure 1B). Further, double knockdown via bacterial mixing of *unc-37(RNAi)* and *lsy-22(RNAi)* in the *pop-1(q645)* background significantly rescues the loss of DTC phenotype (*lacZ(RNAi)*,100% 0 DTCs; n=125 vs *unc-37(RNAi);lsy-22(RNAi)*, 96% 0 DTCs; n=125) (Supplemental Figure 1B). This initial significant, but mild, evidence of Groucho protein functioning in DTC fate encouraged us to continue our analyses using *unc-37/Groucho or lsy-22/AES* nulls.

For our *unc-37/Groucho* null allele, we utilized allele *tm4649* which contains a large out of frame deletion in the WD repeat domain of *unc-37/Groucho* (Supplemental Figure 1C). For our *lsy-22/AES* null allele, we chose an allele with a nonsense mutation at Q33, located at the beginning of the Q domain (Supplemental Figure 1D). In both cases, the premature stop codons are predicted to further result in gene depletion via non-sense mediated mRNA decay. Next, we combined the anatomical DTC marker *qIs56* [P_*lag-2*_::GFP]) with either the *unc-37(tm4649)* null or *lsy-22(ot244)* null allele. Individual Groucho nulls resulted in a significant increase in DTC fate and thus, a portion of the population with 3 or 4 DTCs (wildtype 0% vs *unc-37/Groucho* null 9.38% or *lsy-22/AES* null 9.37%) (Figure 1C/D). Thus, disruption of UNC-37/Groucho and LSY-22/AES function leads to an expansion of WβA-signaled cell fates, indicating Groucho function is required for proper Wnt-signaled ACDs. Since loss of the coactivator SYS-1/β-catenin results in loss of DTC fate, and because loss of POP-1/TCF or the loss of the POP-1/TCF interaction with SYS-1/β-catenin shows the same phenotype (Siegfried et al., 2004), the DTCs are thought to be an activation dominant tissue. Nevertheless, our results indicate a function of Groucho corepressors is required to limit the signaled cell fate. These data suggest WβA target gene expression must not only be adequately increased in the signaled daughter, but also actively repressed in the unsignaled daughter for proper DTC cell fate determination.

Due to our observation that loss of a single corepressor increased DTC fate, we next evaluated the extent to which UNC-37/Groucho and LSY-22/AES function independently to confer repression of POP-1/TCF regulated target genes. To this end, we generated a double corepressor null using CRISPR/Cas9 to edit the *unc-37(tm4349)* strain, adding a stop codon in the same position as the *lsy-22(ot244)* null allele (Q33STOP). The resulting double Groucho homozygote nulls were early larval lethal. Although this does demonstrate the requirement of Groucho proteins for early developmental processes in embryos derived from heterozygous mothers, it unfortunately precluded the possibility of using the double null to assess the effects of Groucho loss on DTC fate.

To determine the effect of double Groucho loss on DTC fate using an alternative method, we subjected both the *unc-37/Groucho* null to *lsy-22(RNAi)* and the *lsy-22/AES* null to *unc-37(RNAi)*. In both instances, loss of the additional Groucho via RNAi increased the penetrance of the ectopic DTC phenotype (Figure 1E/F). Specifically, in the *unc-37/Groucho* null the addition of *lsy-22(RNAi)* increased the population of worms with extra DTCs to 25.3% from 9.4% in the *unc-37/Groucho* null alone. In the *lsy-22/AES* null, we observed a decrease in worms with 1 DTC when subjected to *unc-37(RNAi)*. Additionally, the *lys-22* null showed a decrease in DTC loss when subjected to *pop-1(RNAi)* compared to wildtype (36.9% vs 75%). Since knockdown of the alternate Groucho protein enhances the phenotype of an individual genetic Groucho null, this suggests that UNC-37/Groucho and LSY-22/AES can function both independently and cooperatively as corepressors. Although Groucho function independent of AES is widely accepted, AES function independent of Groucho contrasts the typical model of Groucho-mediated repression via a Groucho and AES heterocomplex (Muhr, Andersson, Persson, Jessell, & Ericson, 2001).

### UNC-37/Groucho and LSY-22/AES are required for proper seam cell fate

Since we determined Groucho corepression is required for proper cell fate in an activation dominant tissue, we wanted to determine if Grouchos are also critical for cell fate the in the repression dominant SCs. The SCs are an adult stem cell that undergo several WβA-signaled ACDs, and one symmetrical cell division, in their lineage. After each ACD, one terminally differentiated hypodermal cell and one pluripotent seam cell result, giving rise to two sets of 16 pluripotent seam cells that run the length of the worm (Figure 2A). Since loss of the coactivator SYS-1/β-catenin does not affect seam cell fate but depletion of POP-1/TCF greatly expands symmetric cell divisions and SC number, the SCs are considered a repression dominant stem cell type ((Banerjee, Chen, Lin, & Slack, 2010) (Figure 2B) and therefore may be sensitive to corepressor loss.

**Figure 2:**
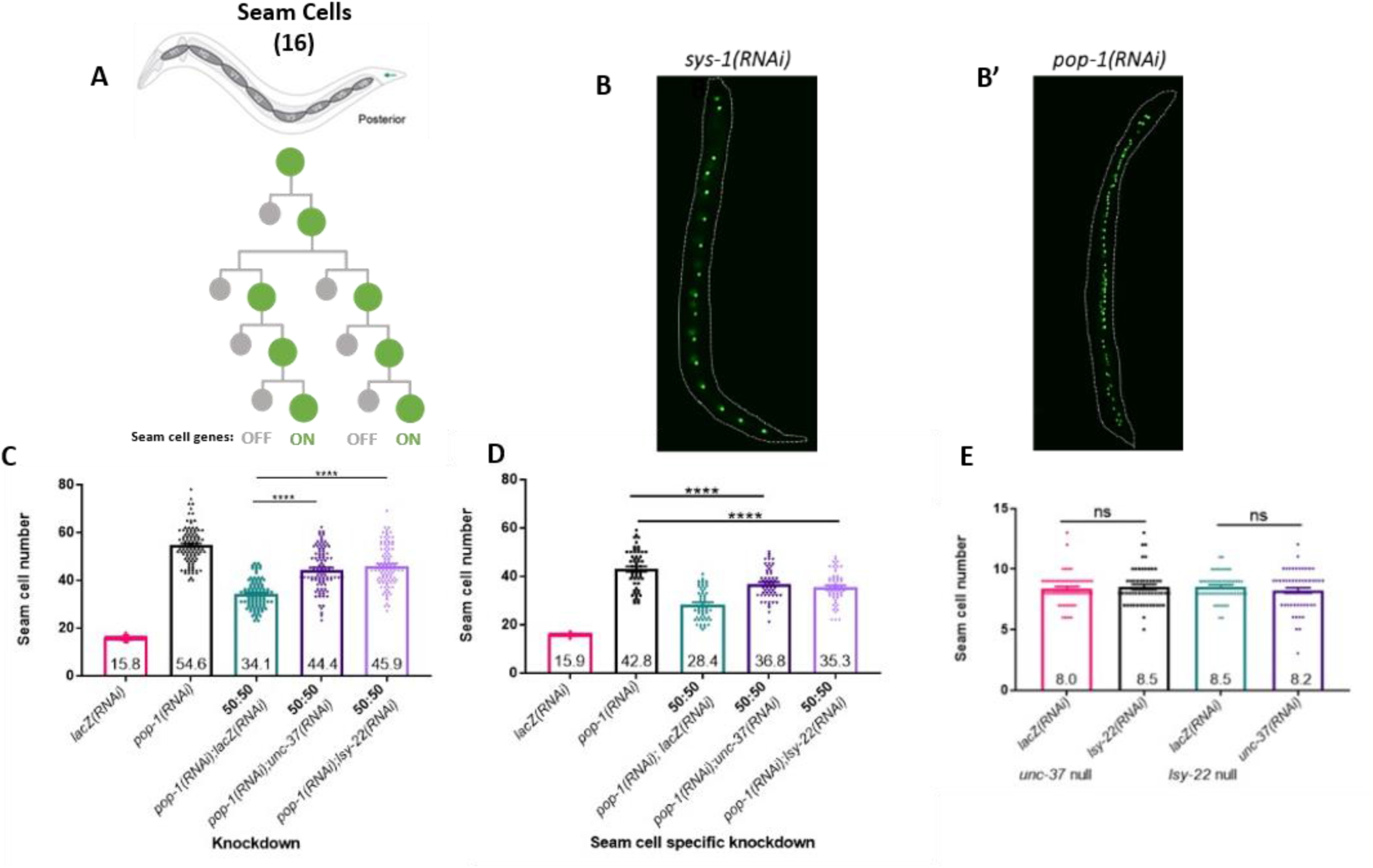
UNC-37/Groucho and LSY-22/AES function in the seam cells. A) Graphic of a representative SC lineage in a wildtype worm. The signaled SC is represented by the green circle and the unsignaled hypodermal cell is represented by the grey circle. B) Representative images showing *sys-1/β-catenin* knockdown does not change SC fate while B’) *pop-1/tcf* knockdown results an expansion of SC fate marked with *wIs51*[P_*scm*_::GFP]. C) Corepressor loss increases SC number in a POP-1/TCF compromised background in whole animal knockdown (n=100, all conditions). D) SC specific knockdown recapitulates full worm knockdown (n=55, all conditions). E) A POP-1/TCF compromised background is required for the effects of Groucho loss in the SCs (n=55, all conditions). Mean value listed inside the bar. *-p≤0.5, **-p<0.01, ***-p<0.001, ****-p≤ 0.0001 via unpaired t-Test.

To determine if Groucho corepressors are required for proper cell fate determination in this repression dominant cell type, we first knocked down *unc-37/Groucho* and *lsy-22/AES* individually and together via RNAi by bacterial feeding. Groucho knockdown alone did not affect cell fate (Supplemental Figure 2A), which may indicate the ability of POP-1/TCF to repress Wnt target genes independent of Groucho corepressors, consistent with the POP-1/TCF depletion phenotype. Since we observed mild suppression of *pop-1(q645)* induced DTC loss by depleting Groucho, we tested the hypothesis that Groucho function in SC divisions could be uncovered in a POP-1/TCF compromised background. To do this, we diluted POP-1/TCF dsRNA by bacterial mixing. First, we utilized an intermediate level of *pop-1/tcf* knockdown by diluting bacteria expressing *pop-1/tcf* dsRNA with an equal amount of bacteria expressing *lacZ* dsRNA (Bekas & Phillips, 2020). Since *lacZ* is a bacterial gene not found in the *C. elegans* genome, it functions as a diluent and is akin to *in vitro* scramble controls. This intermediate level of POP-1/TCF knockdown results in a moderate increase of average SCs to 34.1, which is significantly less than full POP-1/TCF knockdown which averages 54.6 SCs (Figure 2C). Next, instead of diluting *pop-1/tcf* dsRNA with *lacZ* dsRNA we diluted *pop-1/tcf* with bacteria expressing *unc-37/Groucho* or *lsy-22/AES* dsRNA. In both cases, we observed a significant increase of SC number relative to the *pop-1(RNAi);lacZ(RNAi)* phenotype. Specifically, substituting *unc-37(RNAi)* or *lsy-22(RNAi)* for *lacZ(RNAi)* increased the average to 44.5 and 45.9, respectively, from 34.1 (Figure 2C). Notably, the enhancement of seam cell number when *pop-1/tcf* dsRNA is diluted with *unc-37/Groucho* dsRNA or *lsy-22/AES* dsRNA closely recapitulates a full POP-1/TCF knockdown (Figure 2C).

To test if the effects of Groucho loss are increasing SC fate in a cell autonomous manner, we utilized a strain where only the seam cells are susceptible to RNAi. Here, we evaluated a strain where a component of the RISC complex, *rde-1*, is mutated in the entire worm in combination with a wildtype *rde-1* allele provided back under a SC specific promotor (Tabara et al., 1999). This allows for tissue-specific target gene knockdown in the SCs. We subjected this strain to the same Groucho knockdowns in a POP-1/TCF compromised background as above and observed that SC-specific knockdown recapitulated the SC hyperplasia phenotype observed in full worm knockdown (Figure 2D). These results indicate that Groucho functions autonomously in the SCs, rather than any other tissues, to increase SC fate.

As with the DTCs, we next tested the effect of Groucho nulls on seam cell fate. Given the enhancement of SC number observed in RNAi experiments, we anticipated an increase in SC number. However, both the *unc-37/Groucho* null and *lsy-22/AES* null showed a decrease in SC number (average of 8.4 in *unc-37/Groucho* null and average of 8.0 in lsy-22/AES null) (Supplemental Figure 2B/C). Since *unc-37/Groucho* loss has previously been implicated in skipping the singular symmetrical division of this lineage (van der Horst, Cravo, Woollard, Teapal, & van den Heuvel, 2019), we tested if the reduced seam cell number was due to the symmetric division becoming an asymmetric division. To do this, we subjected both *unc-37/Groucho* null and *lsy-22/AES* null to knockdown of a gene, *lin-28*, known to result in a skip of the symmetric division (Vadla, Kemper, Alaimo, Heine, & Moss, 2012). In both cases, *lin-28* knockdown did not enhance the SC loss phenotype observed in the individual Groucho nulls (Supplemental Figure 2D). Thus, we conclude that the decreased number of SCs results from improper cell specification during the single symmetric division the SC linage undergoes. Since we are testing Groucho function in WβA-signaled asymmetric cell divisions, not symmetric divisions, we continued to evaluate the role of Groucho function in the genetic Groucho null lines assuming a new, lower seam cell average of 8 rather than the wildtype seam cell number of 16.

Next, we evaluated the effect of double Groucho loss on SC fate. Since the CRISPR-generated genetic double null is early larval lethal, we again performed RNAi to knockdown the wildtype Groucho allele in each individual Groucho null line. However, we did not see a deviation from the new, adjusted seam cell number of 8 (Figure 2E). We did observe occasional ectopic P_scm_::GFP misexpression in the tail of *lsy-22/AES* nulls subjected to *unc-37(RNAi)* (Supplemental Figure 2E), which may result from aberrant divisions in the T lineage, and were not included in SC number. For this reason, we conclude that a POP-1/TCF compromised background is necessary to uncover Groucho protein function in seam cell fate specification. Thus, the RNAi data best demonstrates that Groucho proteins are required to maintain proper SC fate determination.

### UNC-37/Groucho regulates POP-1/TCF expression in the seam cells

As we began investigating each the effects of individual Groucho null lines on SC fate, we subjected them to *pop-1(RNAi)* to confirm the well-established phenotype of POP-1/TCF depletion resulting in SC increases (Banerjee et al., 2010). When we subjected the *lsy-22/AES* null to *pop-1(RNAi)*, we saw the expected increase in SC fate (Figure 3A). Unexpectedly, when we subjected the *unc-37/Groucho* null to *pop-1(RNAi)* we saw a severe decrease in SC number from 8 to an average of 2 SCs (Figure 3A). To probe the cause of this unforeseen SC decrease, we first tested the possibility of excess *lsy-22/AES* in the *unc-37/Groucho* null contributing to the loss of SC phenotype. Since UNC-37/Groucho and LSY-22/AES are thought to work in a heterocomplex, we tested the possibility that excess LSY-22/AES in the *unc-37/Groucho* null results in SC loss. To this end, we subjected the *unc-37/Groucho* null to double knockdown of *lsy-22(RNAi)* and *pop-1(RNAi)* via bacterial mixing. If the decreased SC phenotype was due to excess LSY-22/AES, we would expect to rescue the decreased SC phenotype. However, double knockdown of *lsy-22/AES* and *pop-1* did not rescue the decreased SC phenotype observed in *unc-37/Groucho* nulls subjected to *pop-1(RNAi)* alone (average 2.1 vs 2.2, respectively) (Supplemental Figure 3A). We therefore conclude the decreased SC number observed is not a result of excess *lsy-22/AES*.

**Figure 3:**
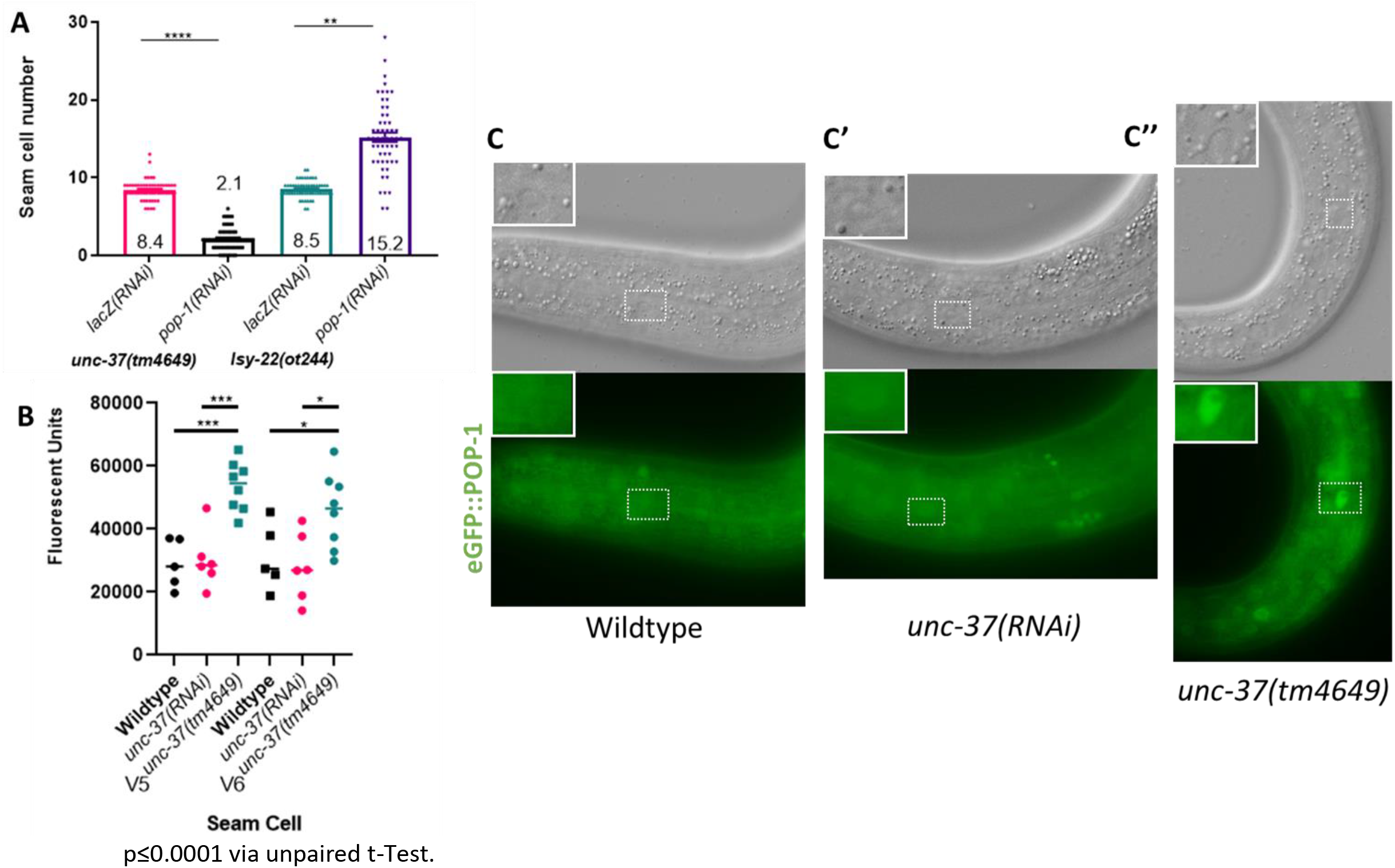
UNC-37/Groucho regulates POP-1/TCF expression in the seam cells. A) *pop-1(RNAi)* decreases SC number in the *unc-37/Groucho* null (mean value listed inside or on top of the bar; n=55, all conditions). B) quantification of *pop-1(he335[eGFP::loxP::pop-1])* in either V5 or V6 SC for wildtype, *unc-37(RNAi)* or, *unc-37/Groucho* null worms. Mean value denoted with straight line (wildtype V5 & V6, n=5; *unc-37(RNAi)* V5 & V6, n=6; *unc-37/Groucho* null V5 & V6, n=8). C) Representative images of wildtype C’) *unc-37(RNAi)* and C’’) *unc-37/Groucho* null SCs expressing *pop-1(he335[eGFP::loxP::pop-1])*. Quantified SC outlined in a white dotted box and enlarged in inset. *-p≤0.5, **-p<0.01, ***-p<0.001, ****-

Since the decrease in SC number is not a result of excess *lsy-22/AES*, we next investigated the extent to which UNC-37/Groucho regulates *pop-1/tcf* expression. Because Grouchos are broadly utilized corepressors, it is possible UNC-37/Groucho represses *pop-1/tcf* in the SCs. If so, interpreting the results from an *unc-37/Groucho* null in the SCs would be complicated by changes to POP-1/TCF levels. To test this, we combined a transgene marking the SC adherens junctions (AJM-1::mcherry) as a cell type marker with an eGFP tagged endogenous *pop-1*(he335[eGFP::loxP::pop-1]). We then quantified the amount of POP-1/TCF protein present in developmentally matched *unc-37/Groucho* nulls relative to *unc-37(RNAi)* and wildtype. The results showed a significant increase in POP-1/TCF protein levels in the *unc-37/Groucho* null relative to *unc-37(RNAi)* and wildtype controls (Figure 3B/C). These data indicate that *unc-37/Groucho* negatively regulates the expression of its own transcription factor *pop-1/tcf* in the SCs, providing an explanation for the decreased SC phenotype in *unc-37(tm4649);pop-1(RNAi)* animals. Moreover, the increase of POP-1/TCF protein in the SCs of the *unc-37/Groucho* null is not observed in *unc-37(RNAi)*. This result furthers the rationale that Groucho corepressor function in the SCs is best assessed in knockdown, rather than knockout, perturbations where we see insignificant change to POP-1/TCF levels.

### Groucho regulation of Wnt target genes is context dependent

The anatomical markers for both the DTCs and SCs fate rely on outputs further downstream from the direct SYS-1/β-catenin interaction with POP-1/TCF at Wnt target genes. For this reason, we wanted to evaluate the direct effect of Groucho loss on Wnt target gene expression in the DTCs and SCs. Here, we utilized fluorescent protein output controlled by the promoter of known DTC and SC Wnt target genes, *ceh-22::venus* and *egl-18a::H1-Wcherry*, respectively. We then subjected these strains to the same Groucho perturbations where cell fate changes occurred to evaluate the effect on Wnt target gene expression.

In the DTCs, in both the single and double Groucho loss conditions, we found significantly more worms exhibited cells derepressing the *ceh-22* target gene than underwent cell fate changes (Figure 4A/B). This was the case for the *unc-37/Groucho* null alone (9.4% *qIs56*[P_lag-2_::GFP] vs 41.9% *qIs90*[P_ceh-22_::YFP]) or with the addition of *lsy-22(RNAi)* (25.3% *qIs56*[P_lag-2_::GFP] vs 70% *qIs90*[P_ceh-22_::YFP]) and for the *lsy-22/AES null* alone (9.4% *qIs56*[P_lag-2_::GFP] vs 20% *qIs90*[P_ceh-22_::YFP]) or with the addition of *unc-37(RNAi)* (9.2% *qIs56*[P_lag-2_::GFP] vs 30% *qIs90*[P_ceh-22_::YFP]) (Figure 4A/B). To confirm the cells misexpressing Wnt target genes were not lingering fluorescent positive cells from previous divisions, we measured gonadal length as a proxy for developmental timing. If the extra *ceh-22* expressing cells were a caveat of developmental timepoint, we would expect gonadal length of worms with ectopic expression to cluster around a similar distance, indicating similar developmental stage. Instead, we found worms with *ceh-22* misexpression were widely dispersed with respect to gonadal length (Supplemental Figure 4A/B/C). Thus, the cells exhibiting derepressed *ceh-22* are not a result of developmental delays in the DTC linage. Taken together, our findings indicate that, in Groucho-depleted larvae, more cells in the somatic gonadal lineage derepress Wnt target genes without becoming a *bona fide* DTC expressing the terminal cell fate marker. This indicates the presence of an intermediate cell type not captured by evaluation of the anatomical DTC marker. Thus, our findings suggest that loss of Groucho function in an activation dominant tissue results in an increase of intermediate cell fates more frequently than cell fate changes.

**Figure 4:**
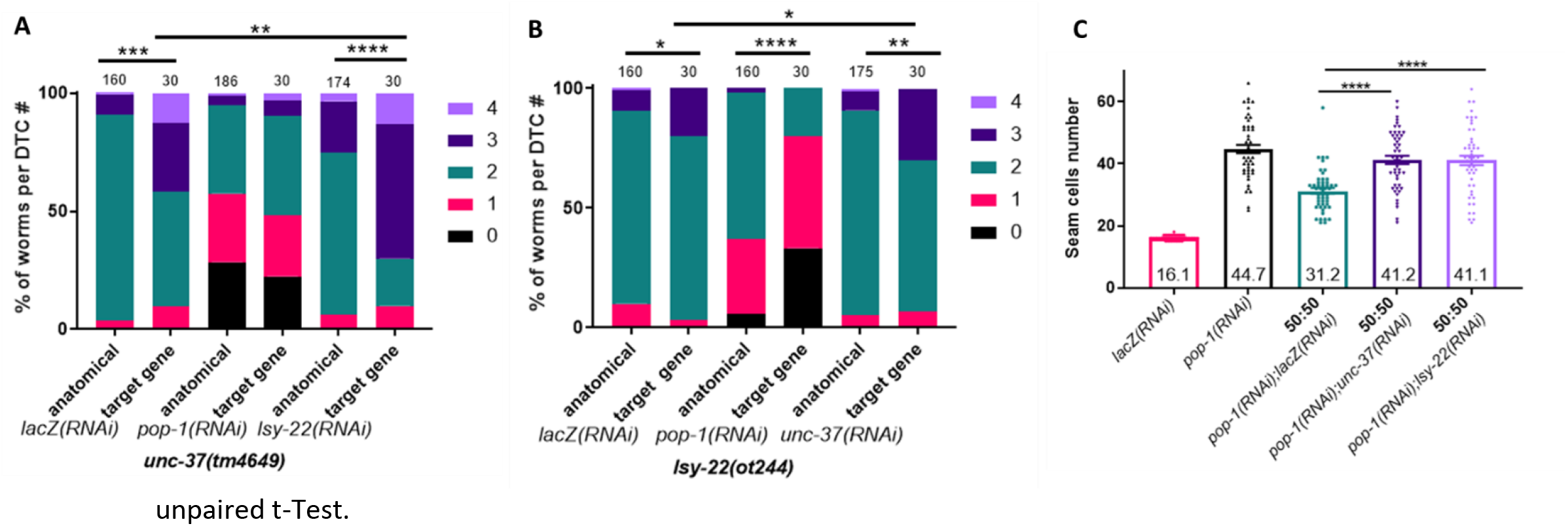
UNC-37/Groucho and LSY-22/AES regulate Wnt target gene expression in the SCs and DTCs. A) DTC target gene derepression and cell fate changes in *unc-37/Groucho* null (n values above each bar). B) DTC target gene derepression and cell fate changes in *lsy-22/AES* null (n values above each bar). C) Corepressor loss increases Wnt target gene expression in a POP-1/TCF compromised background (mean value listed inside the bar; n=50, all conditions). *-p≤0.5, **-p<0.01, ***-p<0.001, ****-p≤0.0001 via

Next, we investigated the effect of Groucho loss on Wnt target gene expression in the SCs. When looking in the *pop-1/tcf* compromised background, we found Wnt target gene derepression recapitulated increases observed in anatomically marked cells (Figure 4C). This result contrasts the observed intermediate cell population in the activation dominant DTC tissue. This indicates a full cell fate conversion occurs from Groucho loss in the repression dominant tissue, where activation dominant tissues with Groucho loss derepress Wnt target genes more frequently than the occurrence of cell fate conversion. Taken together, we observe an unexpected difference in Wnt target gene derepression and cell fate changes with Groucho loss in the activation dominant DTCs and the expected similarity of Wnt target gene derepression and cell fate changes with Groucho loss in the repression dominant SCs.

### Corepressors function in embryonic endoderm specification

To determine the pervasiveness of Groucho regulation in WβA-signaled ACD, we next evaluated their role in embryonic endoderm development. The endoderm develops from ACDs of the EMS cell. The EMS mother cell divides to give rise to the mesoderm (MS) cell and the endoderm (E) cell (Figure 5a). The endoderm cell then undergoes several divisions ultimately giving rise to the intestine while the MS cell linage gives rise to muscle, pharynx, and neuronal cell types. To evaluate Groucho loss induced changes in endoderm development, we first utilized a transgene expressing a histone tagged GFP under the *end-1* gene promotor, *teIs46[P_end-1_::GFP::H2B]*, to assess changes expression changes of the known endodermal WβA target gene. In this strain, the transgene begins expression marking the endoderm linage at the stage with 2E daughter cells (2E), one division after the E and MS specification. Previous studies show loss of the SYS-1/β-catenin coactivator does not affect the number of cells expressing the endoderm transgene, although it does result in dimmer transgene expression in these signaled cells (Figure 5B) (Siegfried et al., 2004). Properly specifying endoderm in the absence of the coactivator categorizes this ACD as repression dominant. This category is also supported by evidence that POP-1/TCF loss results in the two MS cells misexpressing the endoderm marker (Figure 5B) (Maduro & Rothman, 2002). Further, when investigating the effects of *pop-1(RNAi)* on mesodermal target genes, the 2E daughters misexpressed the mesoderm marker, P_tbx-35_::GFP (Owraghi, Broitman-Maduro, Luu, Roberson, & Maduro, 2010). This indicates an intermediate phenotype, where loss of *pop-1/tcf* results in misexpression of mesoderm genes in the E linage and endoderm genes in the MS lineage, similar to our observed effects in the DTC specification (Figure 4A/B).

**Figure 5:**
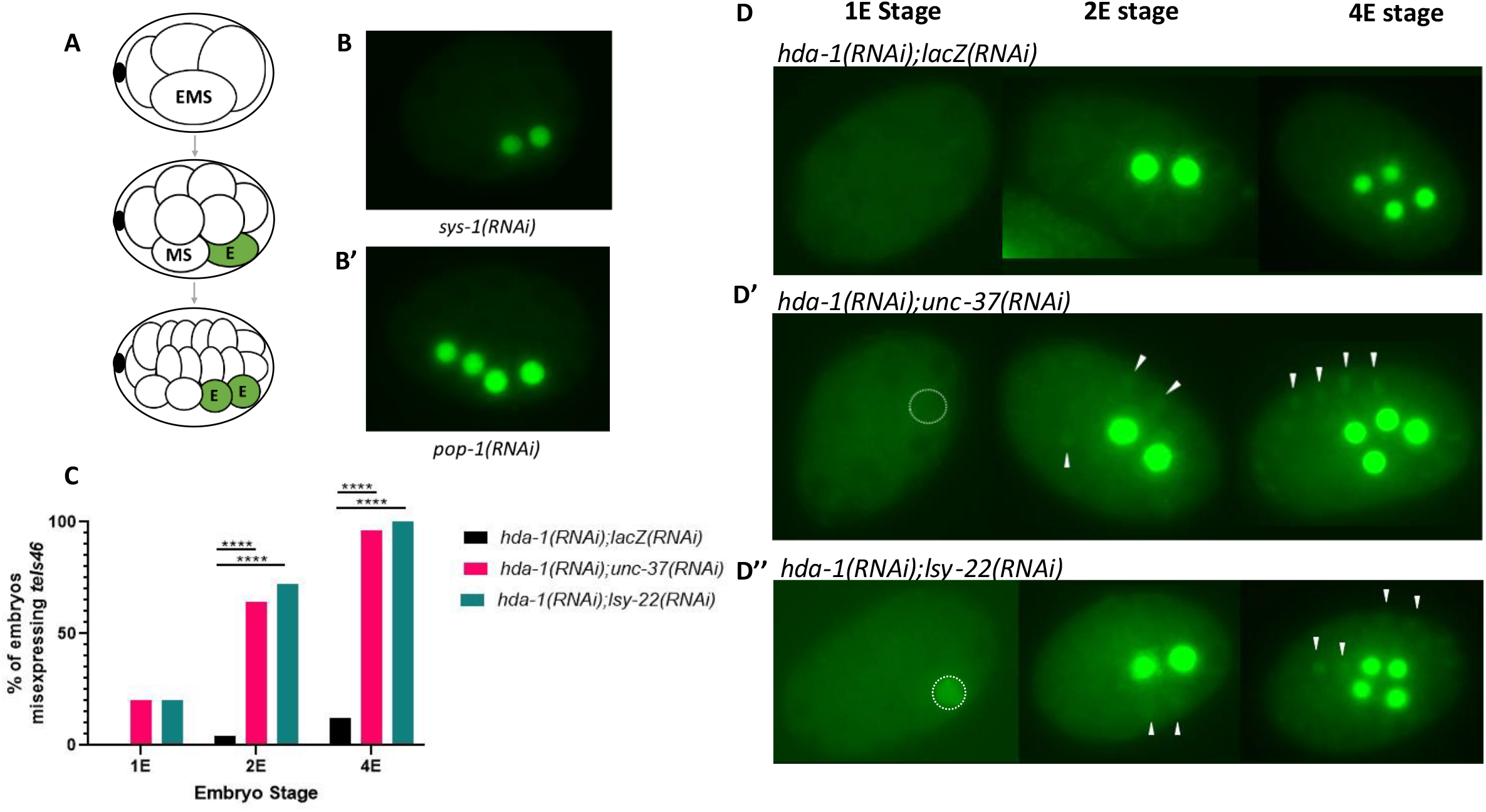
Corepressors function in embryonic endoderm specification. A) Graphic of the endoderm lineage from EMS to 2E stage where the E linage receives the Wnt signal. B) Representative images where *sys-1/β-catenin* knockdown does not affect the number of cells expressing *teIs46*[P_end-1_::GFP::H2B] but B’) *pop-1/tcf* knockdown derepresses *teIs46* [P_end-1_::GFP::H2B] in the mesoderm lineage. C) Percent of population derepressing *teIs46*[P_end-1_::GFP::H2B] (1E, n=10 all conditions; 2E, n=25 all conditions; 4E, n=25 all conditions) when subjected to either control *lacZ(RNAi), unc-37(RNAi)* or *lsy-22(RNAi)* in an *hda-1* compromised background. D) Representative images of *teIs46*[P_end-1_::GFP::H2B] expression at the 1E, 2E and 4E stage resulting from a 50:50 ratio of bacteria containing of *hda-1(RNAi)* with *lacZ(RNAi)* D’) *unc-37(RNAi) or* D”) *lsy-22(RNAi)*. The dotted circle outlines the nuclei of the E cell. White arrow heads denote representative cells ectopically expressing *teIs46*[P_end-1_::GFP::H2B]. *-p≤0.5, **-p<0.01, ***-p<0.001, ****-p≤0.0001 via unpaired t-Test.

To begin our investigation of Groucho function on endoderm development, we subjected worms with the transgene *teIs46[P_end-1_::GFP::H2B]* to knockdown of *unc-37/Groucho, lsy-22/AES*, or a knockdown of both and did not observe any deviation from wildtype (Supplemental Figure 5A), consistent with previous work by Calvo et al 2001. To evaluate the contributions of UNC-37/Groucho independent of LSY-22/AES, and vice versa, we evaluated RNAi of the wildtype Groucho family member in a genetic null of the alternate Groucho, as in the DTCs and SCs. Here, we also had the opportunity to evaluate the CRISPR generated double Groucho null, since a subset of homozygotes are viable through L1. Unexpectedly, we did not see any of changes to cell fate in either the *unc-37/Groucho* null, *lsy-22/AES* null or double *unc-37;lsy-22* null (Supplemental Figure 5B-D). We did not anticipate this result since previous work shows that *unc-37/Groucho* does indeed play a role in endoderm development by enhancing the dsRNA injected phenotype of *hda-1(RNAi)* (Calvo et al., 2001). Since the nulls strains are embryonic lethal and therefore maintained as heterozygotes, it remains possible these corepressors are being maternally inherited in the early embryo, as observed in flies (Delidakis, Preiss, Hartley, & Artavanis-Tsakonas, 1991; Paroush et al., 1994), obscuring our ability to evaluate genetic mutant nulls.

Therefore, to evaluate the possible role for these corepressors in embryonic endoderm development, we turned to evaluating them in a compromised RNAi background. Since *unc-37(RNAi)* enhances the *hda-1(RNAi)* phenotype of *teIs46*[P_end-1_::GFP::H2B] ectopic expression (Calvo et al., 2001), we used this background to validate *unc-37/Groucho* functions in endoderm specification and assess if *lsy-22/AES* functions here as well. We subjected the worms to L1 larval feeding of a 50:50 ratio of either *lacZ(RNAi);hda-1(RNAi), unc-37(RNAi);hda-1(RNAi)* or *lsy-22(RNAi);hda-1(RNAi)* to evaluate changes in transgene expression. In our control *hda-1(RNAi);lacZ(RNAi)*, we rarely found ectopic transgene expression, occurring in 0% of embryos at the 1E stage (n=10), 2% of embryos at the 2E stage (n=25), and 12% of embryos at the 4E stage (n=25) (Figure 5C). When we substituted the control *lacZ* dsRNA for either *unc-37/Groucho* dsRNA or *lsy-22/AES* dsRNA, we observed a significant increase in embryos with ectopic *teIs46*[P_end-1_::GFP::H2B] at the 2E and 4E stages. At the 2E stage, we observed an increase from 2% ectopic expression in our control to 64% or 72% when diluting with either *unc-37/Groucho* (n=25) or *lsy-22/AES* (n=25), respectively (Figure 5C). At the 4E stage we saw an increase of ectopic expression from 12% in our control to 96% or 100% when mixing with *unc-37/Groucho* or *lsy-22/AES*, respectively (n=25 in all conditions) (Figure 5C). Further, in both *unc-37/Groucho* and *lsy-22/AES* knockdown in a *hda-1* compromised background we saw instances of transgene expression at the 1E stage, which is never observed in any controls. Although this occurred at the same frequency for both Groucho knockdowns (20%, n=2/10), it is notable that transgene misexpression a the 1E stage in the *lsy-22(RNAi);hda-1(RNAi)* condition is brighter (Figure 5D). Taken together, the ectopic transgene phenotypes confirm a previously identified role for UNC-37/Groucho function and uncovered a previously unknown role for LSY-22/AES function in repression of endoderm development.

## Discussion

Here, we have shown both *unc-37/Groucho* and *lsy-22/AES* function in cell fate determination of WβA-signaled ACDs. This allows us to update our models of gene regulation in the DTCs, SCs and endoderm determination. In each tissue POP-1/TCF does not function alone to repress Wnt target genes that ultimately determine cell fate but instead POP-1/TCF target gene repression requires a Groucho corepressor.

### Distal tip cell fate requires Groucho and AES

Evaluation of individual *unc-37/Groucho* or *lsy-22/AES* nulls showed an increase in DTC fate, demonstrating a role for Grouch-mediated repression in the activation dominant DTC lineage (Figure 1D). Our results exhibiting *unc-37/Groucho* null enhancement of DTC gains by *lsy-22(RNAi)*, and vice versa, suggests the function of a homo-corepressive complex of *lsy-22/AES* that functions independently of *unc-37/Groucho*, and contrariwise. Thus, we propose updating the model of normal DTC ACD such that the nucleus of the unsignaled daughter contains corepressive complexes composed of *UNC-37/Groucho, LSY-22/AES* or both Groucho family members (Figure 6A). While Gro/TLE has long been thought to function independently of AES (Turki-Judeh & Courey, 2012), this result is the first evidence of a possible *lsy-22/AES* corepressive homocomplex and provides rationale to further investigate other contexts where AES proteins function without Groucho. Surprisingly, in this activation dominant tissue we uncovered a role for corepressors without needing a compromised background; individual *unc-37/Groucho* and *lsy-22/AES* nulls increase DTC fate (9.38% or 9.37%, respectively) which was enhanced to 25.3% in the *unc-37/Groucho* null with *lsy-22(RNAi)*. This may indicate, in activation dominant tissues, Groucho/AES mediated repression may be the predominant repressive mechanism ensuring proper cell fate. Specifically, there is potential for a complex that includes both UNC-37/Groucho and LSY-22/AES as well as UNC-37/Groucho and LSY-22/AES homocomplexes mediating repression in the DTCs.

**Figure 6:**
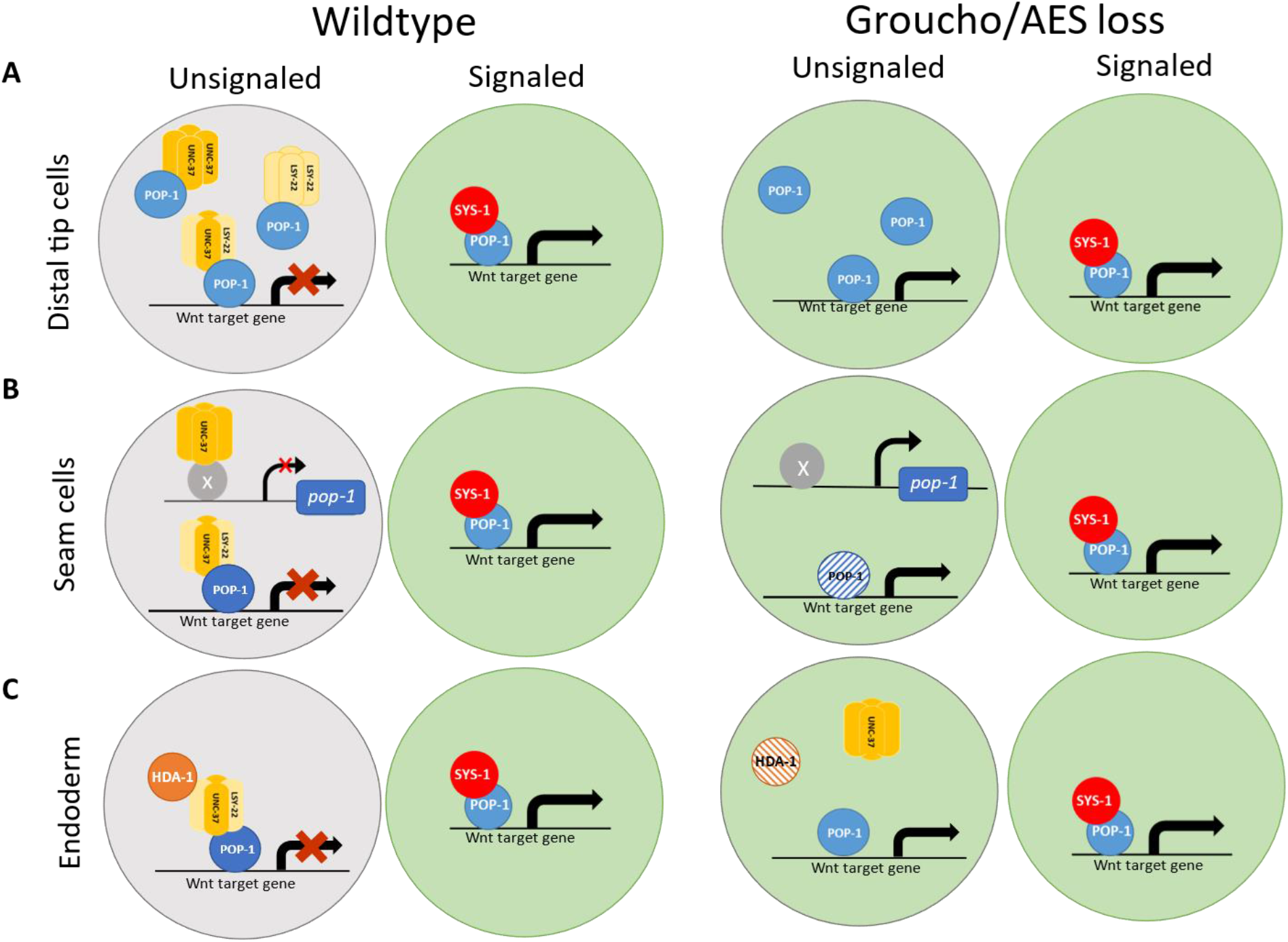
Graphic summary of proposed Groucho function in wildtype and mutant cells. A) Wildtype representation of the DTC nuclei resulting from a Wnt polarized ACD. In the signaled daughter’s nucleus (represented by the green circle) Wnt target genes are transcribed. In wildtype, the unsignaled daughter represses Wnt target genes via interactions with Groucho corepressors (represented by the gray circle). With Groucho and AES loss, the unsignaled daughter derepressed Wnt target genes. B) Wildtype representation of the signaled SC nucleus resulting in POP-1/TCF and SYS-1/β-catenin target gene activation. In the unsignaled daughter, UNC-37/Groucho regulates *pop-1/tcf* and both UNC-37/Groucho and LSY-22/AES work with POP-1/TCF to repress Wnt target gene expression. When subjected to Groucho loss in a POP-1/TCF compromised background (represented by the partially colored POP-1/TCF protein) Wnt target genes are derepressed in the unsignaled cell. C) Wildtype representation of Wnt target gene activation in the signaled endodermal daughter’s nucleus. In the unsignaled nucleus, UNC-37/Groucho and LSY-22/AES work together with HDA-1 to repress Wnt target gene transcription. Loss of LSY-22/AES in an HDA-1 compromised background (represented by the partially colored HDA-1 protein), results in Wnt target gene derepression in the unsignaled daughter’s nucleus.

Further, investigation of Groucho-mediated repression on Wnt target genes demonstrated target gene derepression occurs more frequently than cell fate changes. As such, we identified and intermediate cell type not captured by the distal tip cell fate marker, P_lag-2_::GFP. This demonstrates the importance of Groucho-mediated repression of Wnt target genes in the activation dominant DTCs.

### Seam cell fate requires Groucho and AES

In the SCs, a POP-1/TCF compromised background was needed to uncover the role of Groucho proteins in Wnt target gene regulation and cell fate decisions (Figure 6B). In contrast to the DTCs, Wnt target gene derepression correlated with cell fate changes. Thus, an intermediate cell type was identified in the activation dominant DTCs but not the repression dominant DTCs. Since repression contributes more than activation to SC fate, it is possible multiple mechanisms of repression are working together to ensure proper cell fate. Further, a decrease in SCs was observed in both the *unc-37/Groucho* and *lsy-22/AES* null, indicating these corepressors also play a role in the symmetric division of this lineage. While this has been investigated for *unc-37/Groucho* (van der Horst et al., 2019), a role for *lsy-22/AES* has not previously been implicated in this division. We also uncovered the first instance of *unc-37/Groucho* regulation of *pop-1/TCF*, which speaks to the ubiquity Groucho-mediated repression. Lastly, the closest condition to a double Groucho loss we could evaluate (*unc-37/Groucho* null with *lsy-22(RNAi)* and *lsy-22/AES* null with *unc-37(RNAi)*) did not yield an increased SC phenotype. This indicates either 1) the ability of POP-1/TCF to function as a repressor on its own in repression dominant tissues or 2) POP-1/TCF repression occurring via a Groucho-independent mechanisms. The observation that a *pop-1/tcf* sensitized background was required to uncover Groucho and AES function in SC division is consistent with both possibilities since either scenario utilizes POP-1/TCF in alternative repression mechanisms. Future studies are needed to better understand the full scope of target gene repression in the repression dominant WβA singled ACDs.

### Embryonic endoderm specification requires Groucho and AES

In embryonic endoderm development, our results suggest *UNC-37/Groucho* and *LSY-22/AES* are maternally inherited in the early embryo since RNAi but not homozygous mutants derived from heterozygotes exhibited phenotypes. By knocking down a histone deacetylase known to uncover Groucho function (Calvo et al., 2001), we validated *unc-37/Groucho* function in endoderm specification and demonstrate a novel role for *lsy-22/AES* in this repression dominant cell fate decision (Figure 6C). Similar mechanisms of repression may be utilized in both the SCs and endoderm specification since a compromised background was needed in both repression dominant tissues to uncover Groucho contribution to cell fate.

Creating a compromised background by weakening histone deacetylases provides solid evidence of the Groucho-mediated method of repression in endoderm specification. This does not, however, exclude the possibility that repression is also occurring via interactions with transcriptional machinery or rule out the possibility of Groucho proteins functioning via multiple histone deacetylases. Now that we have shown Groucho and AES proteins function in WβA-signaled ACDs, further studies are needed to continue to uncover the mechanism of repression. Does the method of repression depend on the composition of the Groucho corepressor complex? How universal is an AES only corepressor complex? Is AES-mediated repression only occurring with TCF? How common is AES regulation of other transcription factors that bind the Q domain? These questions all warrant further investigation given this newfound function of Groucho proteins, particularly LSY-22/AES, in *C. elegans* Wnt-signaled ACD.

In summary, our findings have verified LSY-22/AES functions as a *bona fide* corepressor that functions in TCF mediated repression in the nascent unsignaled daughter cell. Interestingly, our work demonstrated the largest contribution of Groucho-mediated repression in the activation dominant DTCs rather than the repression dominant SCs or EMS divisions. Altogether, our studies have demonstrated Groucho-mediated repression occurs in the WβA-signaled divisions of the *C. elegans* DTCs, SCs, and embryonic endoderm.

## Acknowledgments

We thank Josh Thompson and Maria Valdes Michel for helpful comments on the manuscript. Some strains were provided by the Caenorhabditis Genetics Center, which is funded by the National Institute of Health (NIH) Office of Research Infrastructure Programs [P40 OD01440]. We thank Sander van den Heuvel for the strain SV2114 and Sarit Smolikove for our a Cas9 aliquot. This work was supported by NIH GM114007 (BTP), the Predoctoral Training Program in Genetics T32 training grant (T32 GM 008629), the University of Iowa’s Graduate Diversity Fellowship and Ballard-Seashore Dissertation Fellowship.

## Notes

### Competing Interest Statement

The authors have declared no competing interest.

### Summary of Updates

Clarifying redundant Groucho functions nematode/human gene terminology

## References

Bajoghli, B., Aghaallaei, N., Soroldoni, D., & Czerny, T. (2007). The roles of Groucho/Tle in left-right asymmetry and Kupffer’s vesicle organogenesis. Dev Biol, 303(1), 347–361. doi:10.1016/j.ydbio.2006.11.020

Banerjee, D., Chen, X., Lin, S. Y., & Slack, F. J. (2010). kin-19/casein kinase Ialpha has dual functions in regulating asymmetric division and terminal differentiation in C. elegans epidermal stem cells. Cell Cycle, 9(23), 4748–4765. doi:10.4161/cc.9.23.14092

Beagle, B., & Johnson, G. V. (2010). AES/GRG5: more than just a dominant-negative TLE/GRG family member. Dev Dyn, 239(11), 2795–2805. doi:10.1002/dvdy.22439

Bekas, K. N., & Phillips, B. T. (2020). Generating reliable hypomorphic phenocopies by RNAi using long dsRNA as diluent. MicroPubl Biol, 2020. doi:10.17912/micropub.biology.000269

Berk, A. J. (1999). Activation of RNA polymerase II transcription. Curr Opin Cell Biol, 11(3), 330–335. doi:10.1016/S0955-0674(99)80045-3

Blaydon, D. C., Ishii, Y., O’Toole, E. A., Unsworth, H. C., Teh, M. T., Ruschendorf, F., … Kelsell, D. P. (2006). The gene encoding R-spondin 4 (RSPO4), a secreted protein implicated in Wnt signaling, is mutated in inherited anonychia. Nat Genet, 38(11), 1245–1247. doi:10.1038/ng1883

Brenner, S. (1974). The genetics of Caenorhabditis elegans. Genetics, 77(1), 71–94. doi:10.1093/genetics/77.1.71

Calvo, D., Victor, M., Gay, F., Sui, G., Luke, M. P., Dufourcq, P., … Shi, Y. (2001). A POP-1 repressor complex restricts inappropriate cell type-specific gene transcription during Caenorhabditis elegans embryogenesis. EMBO J, 20(24), 7197–7208. doi:10.1093/emboj/20.24.7197

Chen, G., Nguyen, P. H., & Courey, A. J. (1998). A role for Groucho tetramerization in transcriptional repression. Mol Cell Biol, 18(12), 7259–7268. doi:10.1128/MCB.18.12.7259

Conklin, E. G. (1905). The Mutation Theory From the Standpoint of Cytology. Science, 21(536), 525–529. doi:10.1126/science.21.536.525

Daniels, D. L., & Weis, W. I. (2005). Beta-catenin directly displaces Groucho/TLE repressors from Tcf/Lef in Wnt-mediated transcription activation. Nat Struct Mol Biol, 12(4), 364–371. doi:10.1038/nsmb912

Delidakis, C., Preiss, A., Hartley, D. A., & Artavanis-Tsakonas, S. (1991). Two genetically and molecularly distinct functions involved in early neurogenesis reside within the Enhancer of split locus of Drosophila melanogaster. Genetics, 129(3), 803–823. doi:10.1093/genetics/129.3.803

Flowers, E. B., Poole, R. J., Tursun, B., Bashllari, E., Pe’er, I., & Hobert, O. (2010). The Groucho ortholog UNC-37 interacts with the short Groucho-like protein LSY-22 to control developmental decisions in C. elegans. Development, 137(11), 1799–1805. doi:10.1242/dev.046219

He, X., Semenov, M., Tamai, K., & Zeng, X. (2004). LDL receptor-related proteins 5 and 6 in Wnt/beta-catenin signaling: arrows point the way. Development, 131(8), 1663–1677. doi:10.1242/dev.01117

Hefel, A., & Smolikove, S. (2019). Tissue-Specific Split sfGFP System for Streamlined Expression of GFP Tagged Proteins in the Caenorhabditis elegans Germline. G3 (Bethesda), 9(6), 1933–1943. doi:10.1534/g3.119.400162

Horvitz, H. R., & Herskowitz, I. (1992). Mechanisms of asymmetric cell division: two Bs or not two Bs, that is the question. Cell, 68(2), 237–255. doi:10.1016/0092-8674(92)90468-r

Kimelman, D., & Xu, W. (2006). beta-catenin destruction complex: insights and questions from a structural perspective. Oncogene, 25(57), 7482–7491. doi:10.1038/sj.onc.1210055

Kornberg, R. D. (2005). Mediator and the mechanism of transcriptional activation. Trends Biochem Sci, 30(5), 235–239. doi:10.1016/j.tibs.2005.03.011

Li, S. S. (2000). Structure and function of the Groucho gene family and encoded transcriptional corepressor proteins from human, mouse, rat, Xenopus, Drosophila and nematode. Proc Natl Sci Counc Repub China B, 24(2), 47–55. Retrieved from https://www.ncbi.nlm.nih.gov/pubmed/10809080

Logan, C. Y., & Nusse, R. (2004). The Wnt signaling pathway in development and disease. Annu Rev Cell Dev Biol, 20, 781–810. doi:10.1146/annurev.cellbio.20.010403.113126

Maduro, M. F., & Rothman, J. H. (2002). Making worm guts: the gene regulatory network of the Caenorhabditis elegans endoderm. Dev Biol, 246(1), 68–85. doi:10.1006/dbio.2002.0655

Malik, S., & Roeder, R. G. (2010). The metazoan Mediator co-activator complex as an integrative hub for transcriptional regulation. Nat Rev Genet, 11(11), 761–772. doi:10.1038/nrg2901

Miyasaka, H., Choudhury, B. K., Hou, E. W., & Li, S. S. (1993). Molecular cloning and expression of mouse and human cDNA encoding AES and ESG proteins with strong similarity to Drosophila enhancer of split groucho protein. Eur J Biochem, 216(1), 343–352. doi:10.1111/j.1432-1033.1993.tb18151.x

Muhr, J., Andersson, E., Persson, M., Jessell, T. M., & Ericson, J. (2001). Groucho-mediated transcriptional repression establishes progenitor cell pattern and neuronal fate in the ventral neural tube. Cell, 104(6), 861–873. doi:10.1016/s0092-8674(01)00283-5

Neumuller, R. A., & Knoblich, J. A. (2009). Dividing cellular asymmetry: asymmetric cell division and its implications for stem cells and cancer. Genes Dev, 23(23), 2675–2699. doi:10.1101/gad.1850809

Owraghi, M., Broitman-Maduro, G., Luu, T., Roberson, H., & Maduro, M. F. (2010). Roles of the Wnt effector POP-1/TCF in the C. elegans endomesoderm specification gene network. Dev Biol, 340(2), 209–221. doi:10.1016/j.ydbio.2009.09.042

Paix, A., Schmidt, H., & Seydoux, G. (2016). Cas9-assisted recombineering in C. elegans: genome editing using in vivo assembly of linear DNAs. Nucleic Acids Res, 44(15), e128. doi:10.1093/nar/gkw502

Parma, P., Radi, O., Vidal, V., Chaboissier, M. C., Dellambra, E., Valentini, S., … Camerino, G. (2006). R-spondin1 is essential in sex determination, skin differentiation and malignancy. Nat Genet, 38(11), 1304–1309. doi:10.1038/ng1907

Paroush, Z., Finley, R. L., Jr., Kidd, T., Wainwright, S. M., Ingham, P. W., Brent, R., & Ish-Horowicz, D. (1994). Groucho is required for Drosophila neurogenesis, segmentation, and sex determination and interacts directly with hairy-related bHLH proteins. Cell, 79(5), 805–815. doi:10.1016/0092-8674(94)90070-1

Pflugrad, A., Meir, J. Y., Barnes, T. M., & Miller, D. M., 3rd. (1997). The Groucho-like transcription factor UNC-37 functions with the neural specificity gene unc-4 to govern motor neuron identity in C. elegans. Development, 124(9), 1699–1709. Retrieved from https://www.ncbi.nlm.nih.gov/pubmed/9165118

Phillips, B. T., Kidd, A. R., 3rd, King, R., Hardin, J., & Kimble, J. (2007). Reciprocal asymmetry of SYS-1/beta-catenin and POP-1/TCF controls asymmetric divisions in Caenorhabditis elegans. Proc Natl Acad Sci U S A, 104(9), 3231–3236. doi:10.1073/pnas.0611507104

Pinto, M., & Lobe, C. G. (1996). Products of the grg (Groucho-related gene) family can dimerize through the amino-terminal Q domain. J Biol Chem, 271(51), 33026–33031. doi:10.1074/jbc.271.51.33026

Rhyu, M. S., Jan, L. Y., & Jan, Y. N. (1994). Asymmetric distribution of numb protein during division of the sensory organ precursor cell confers distinct fates to daughter cells. Cell, 76(3), 477–491. doi:10.1016/0092-8674(94)90112-0

Schneider, S. Q., & Bowerman, B. (2007). beta-Catenin asymmetries after all animal/vegetal-oriented cell divisions in Platynereis dumerilii embryos mediate binary cell-fate specification. Dev Cell, 13(1), 73–86. doi:10.1016/j.devcel.2007.05.002

Siegfried, K. R., Kidd, A. R., 3rd, Chesney, M. A., & Kimble, J. (2004). The sys-1 and sys-3 genes cooperate with Wnt signaling to establish the proximal-distal axis of the Caenorhabditis elegans gonad. Genetics, 166(1), 171–186. doi:10.1534/genetics.166.1.171

Song, H., Hasson, P., Paroush, Z., & Courey, A. J. (2004). Groucho oligomerization is required for repression in vivo. Mol Cell Biol, 24(10), 4341–4350. doi:10.1128/MCB.24.10.4341-4350.2004

Tabara, H., Sarkissian, M., Kelly, W. G., Fleenor, J., Grishok, A., Timmons, L., … Mello, C. C. (1999). The rde-1 gene, RNA interference, and transposon silencing in C. elegans. Cell, 99(2), 123–132. doi:10.1016/s0092-8674(00)81644-x

Timmons, L., & Fire, A. (1998). Specific interference by ingested dsRNA. Nature, 395(6705), 854. doi:10.1038/27579

Turki-Judeh, W., & Courey, A. J. (2012). Groucho: a corepressor with instructive roles in development. Curr Top Dev Biol, 98, 65–96. doi:10.1016/B978-0-12-386499-4.00003-3

Vadla, B., Kemper, K., Alaimo, J., Heine, C., & Moss, E. G. (2012). lin-28 controls the succession of cell fate choices via two distinct activities. PLoS Genet, 8(3), e1002588. doi:10.1371/journal.pgen.1002588

van der Horst, S. E. M., Cravo, J., Woollard, A., Teapal, J., & van den Heuvel, S. (2019). C. elegans Runx/CBFbeta suppresses POP-1 TCF to convert asymmetric to proliferative division of stem cell-like seam cells. Development, 146(22). doi:10.1242/dev.180034

Winnier, A. R., Meir, J. Y., Ross, J. M., Tavernarakis, N., Driscoll, M., Ishihara, T., … Miller, D. M., 3rd. (1999). UNC-4/UNC-37-dependent repression of motor neuron-specific genes controls synaptic choice in Caenorhabditis elegans. Genes Dev, 13(21), 2774–2786. doi:10.1101/gad.13.21.2774

Zhang, H., & Emmons, S. W. (2002). Caenorhabditis elegans unc-37/groucho interacts genetically with components of the transcriptional mediator complex. Genetics, 160(2), 799–803. doi:10.1093/genetics/160.2.799

