## Supplemental data for "*unc-37/*Groucho and *lsy-22/*AES repress Wnt target genes in *C. elegans* asymmetric cell divisions"

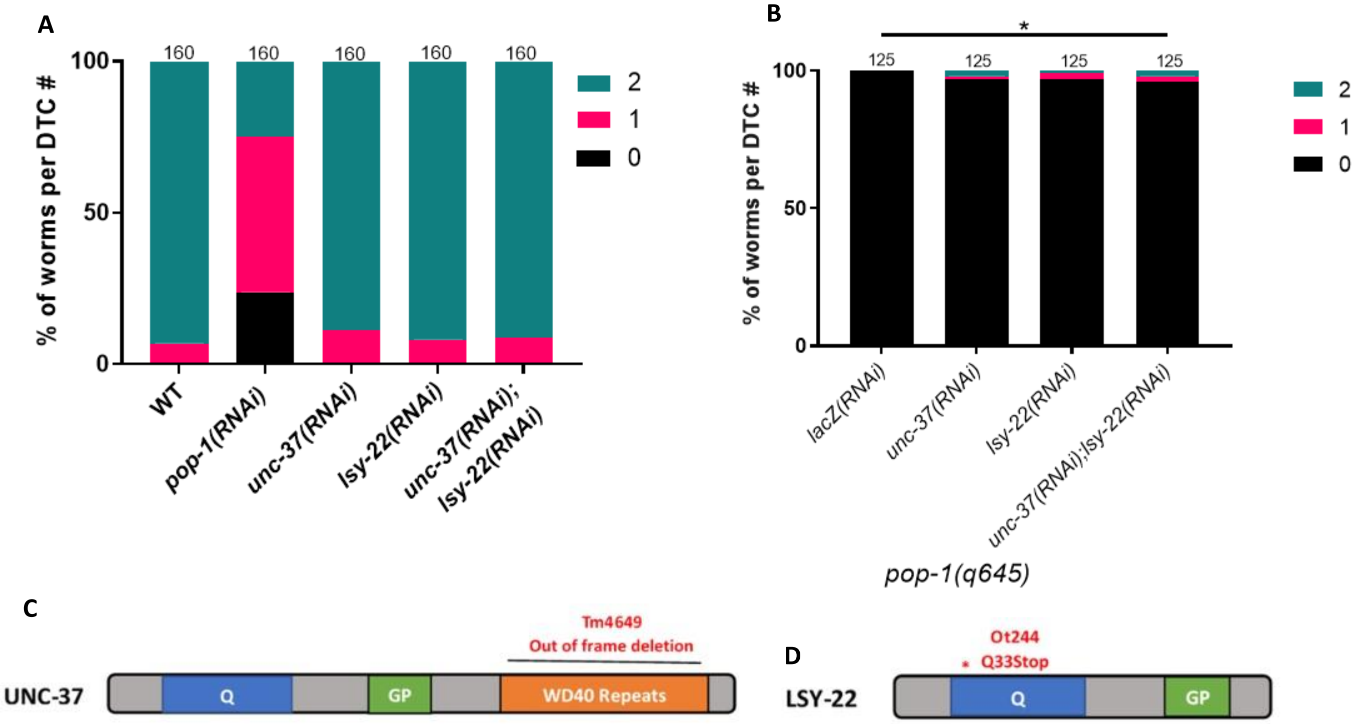


**Supplemental Figure 1: Evaluating Groucho knockdown and graphics of null alleles.** A) Corepressor knockdown does not change DTC fate. B) Double Groucho knockdown partially suppressed DTC loss from defective POP-1/TCF activation allele. C) Graphic of *unc-37/Groucho* null allele *tm4649* D) Graphic of *lsy-22/AES* null allele *ot244.* N value listed above each bar. *-p≤0.5, **-p<0.01, ***-p< 0.001, ****-p≤ 0.0001 via unpaired t-Test.

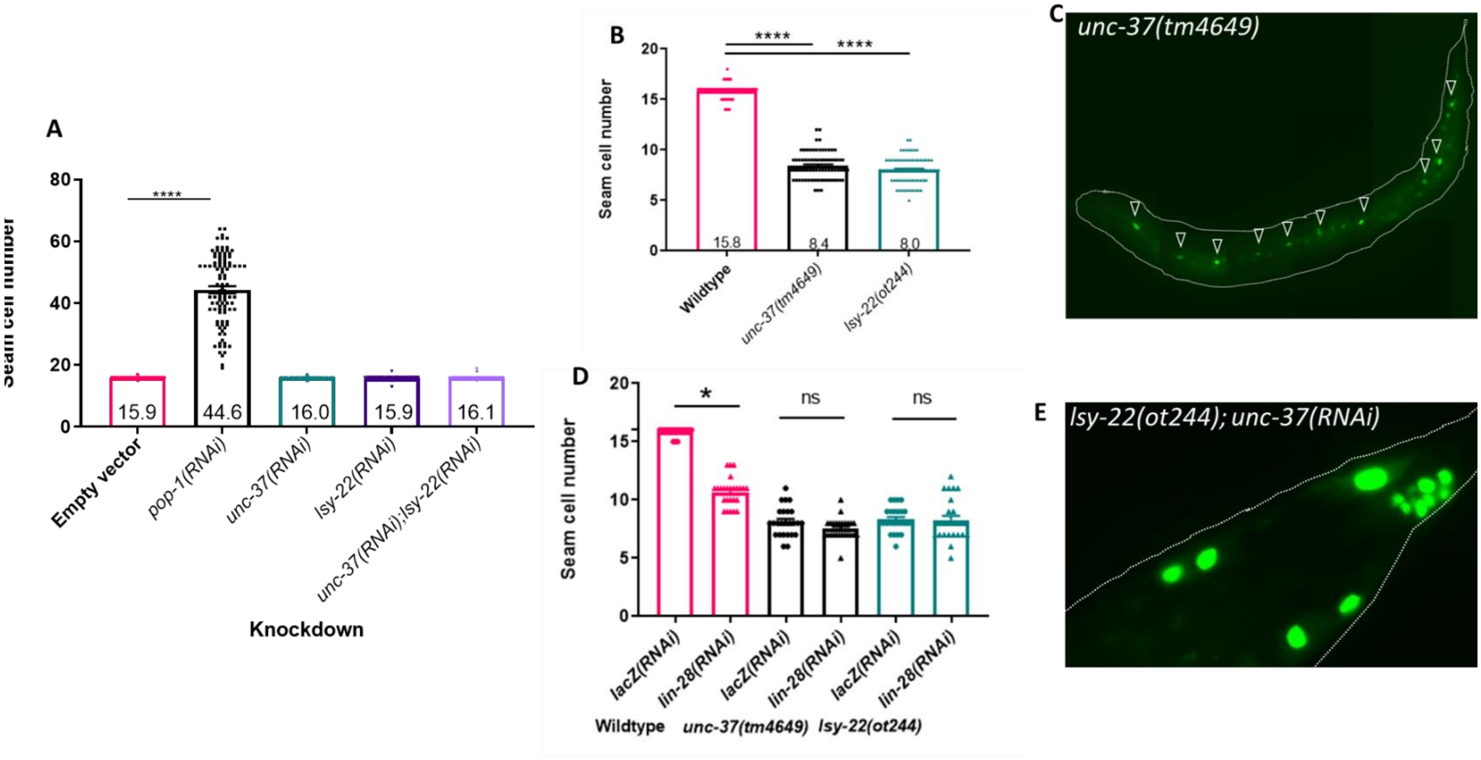


**Supplemental Figure 2: Groucho nulls, but not knockdown, skip the symmetric division and misspecify tail SC lineage.** A) Corepressor knockdown does not affect SC fate (mean value listed inside the bar; n=100 all conditions). B) SC fate decreases in *unc-37/Groucho* and *lsy-22/AES* nulls (mean value listed inside the bar; n=75 all conditions). C) Representative image of decreased SCs in the *unc-37/Groucho* null. Arrow heads denote SCs. D) Loss of symmetric division does not affect SC fate in Groucho nulls (n=25, all conditions). E) Representative image of tail cells misexpressing $P_{scm}$::GFP in the *lsy-22/AES* null with *unc-37(RNAi).* *-p≤0.5, **-p<0.01, ***-p< 0.001, ****-p≤ 0.0001 via unpaired t-Test.

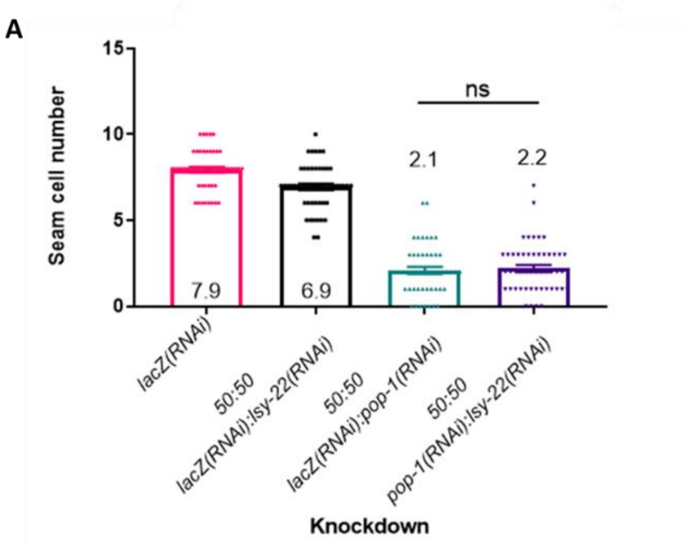


**Supplemental Figure 3: Decreased SC number in *unc-37/Groucho* null with *pop-1/tcf* knockdown is not a function of excess LSY-22/AES.** A) *lsy-22/AES* knockdown does not rescue decreased SC phenotype in *unc-37/Groucho* null with *pop-1/tcf* knockdown (N=55, for all conditions)

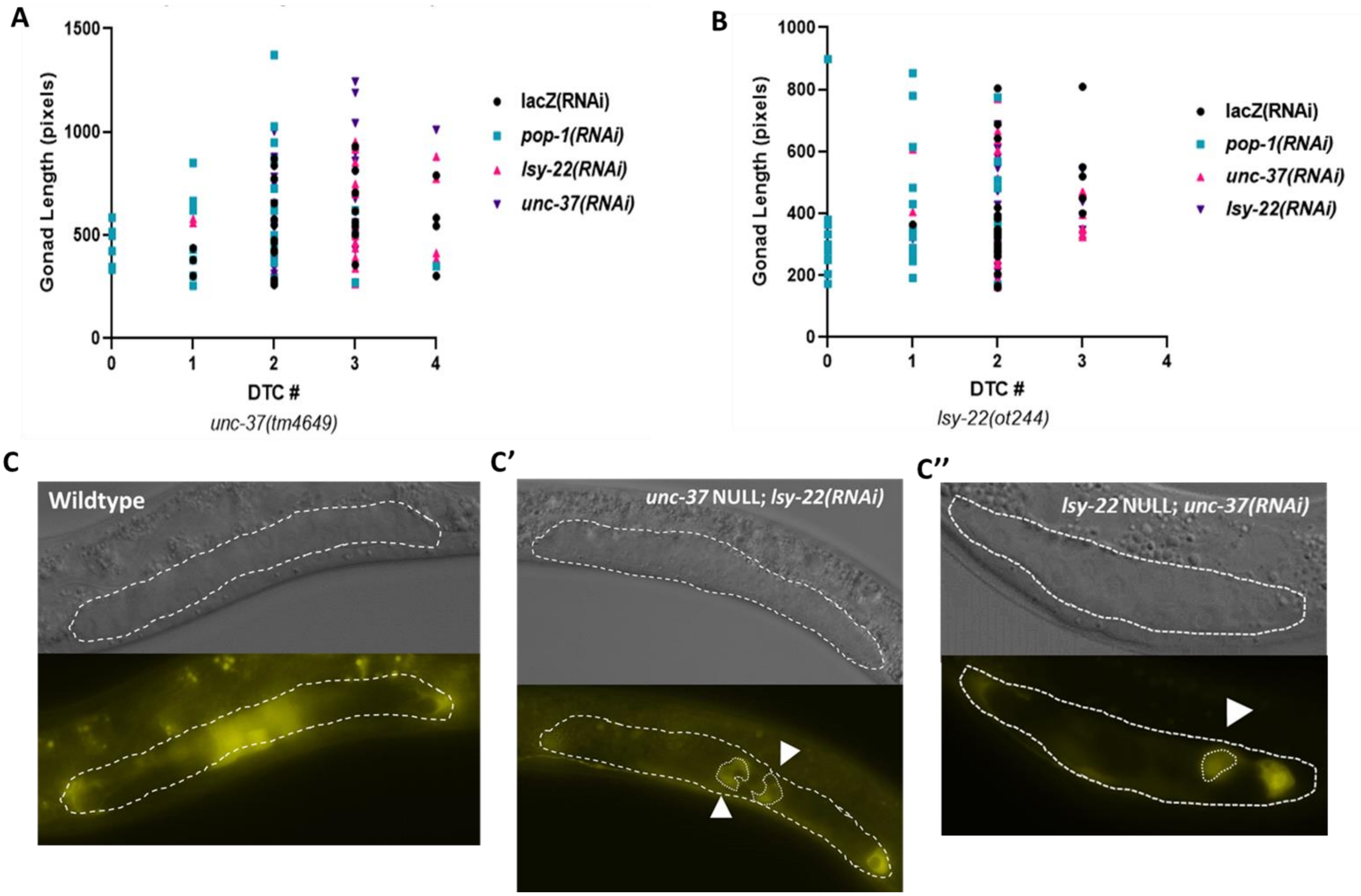


**Supplemental Figure 4: Wnt target gene expression is independent of gonadal development.** A) *qIs90*[$P_{ceh\text{-}22}$::YFP] misexpression in *unc-37/Groucho* null and RNAi conditions does not cluster around one gonadal developmental timepoint (n=30, all conditions). B) *qIs90*[$P_{ceh\text{-}22}$::YFP] misexpression in *lsy-22/AES* null and RNAi conditions do not cluster around one gonadal developmental timepoint (n=30, all conditions). C) Representative images of wildtype *qIs90*[$P_{ceh\text{-}22}$::YFP] expression and misexpression in C’) *unc-37/Groucho* null with *lsy-22(RNAi)* or C’’) *lsy-22/AES* null with *unc-37(RNAi)*. Here, the gonad is outlined with white dashed lines and cells ectopically expressing *qIs90*[$P_{ceh\text{-}22}$::YFP] are outlined with a white dotted line and denoted by white arrowheads.

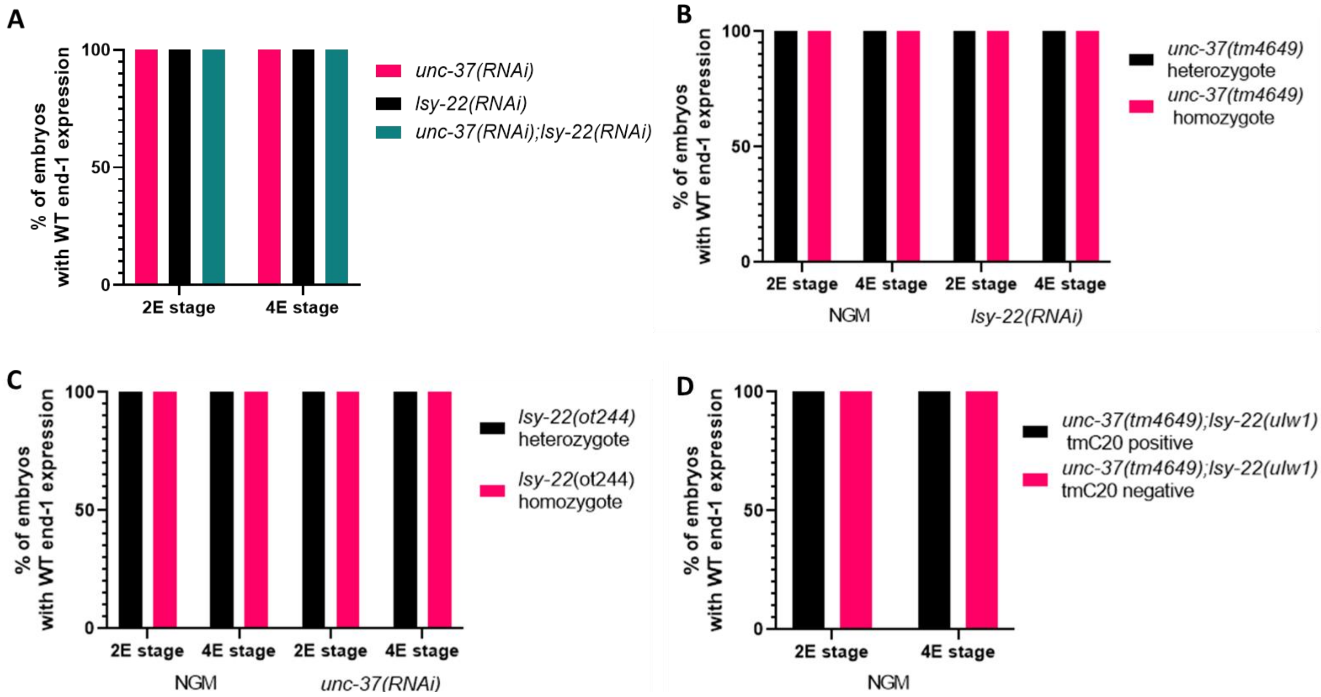


**Supplemental Figure 5: Corepressors function in embryonic endoderm specification Groucho proteins are maternally inherited in the early embryo** A) Groucho knockdown does not affect endoderm fate from *teIs46*[$P_{end-1}$::GFP::H2B](n=20, all conditions). B) *unc-37/Groucho* null does not affect endoderm development with *teIs46*[$P_{end-1}$::GFP::H2B](heterozygotes NGM, n=12; homozygotes NGM n=5; heterozygotes *lsy-22(RNAi)* n=10; homozygotes *lsy-22(RNAi)* n=5). C) *lsy-22/AES* null does not affect endoderm development with *teIs46*[$P_{end-1}$::GFP::H2B](heterozygotes NGM, n=14; homozygotes NGM n=5; heterozygotes *lsy-22(RNAi)* n=10; homozygotes *lsy-22(RNAi)* n=5). D) Double *unc-37/Groucho, lsy-22/AES* null does not affect endoderm development with *teIs46*[$P_{end-1}$::GFP::H2B](tmC20 positive n=10; tmC20 negative n=5).
